# Species distribution modeling of the castor bean tick, *Ixodes ricinus* (Linnaeus, 1758), under current and future climates, with a special focus on Ukraine and Latvia

**DOI:** 10.1101/2024.07.03.602006

**Authors:** Volodymyr Tytar, Iryna Kozynenko, Mihails Pupins, Arturs Škute, Andris Čeirāns, Oksana Nekrasova

## Abstract

This study assesses the impact of climate change on the distribution of *Ixodes ricinus* ticks, that is transmitting Lyme disease, a growing public health concern. Utilizing ensemble models from the R package flexsdm and climate data from WorldClim, ENVIREM, and CliMond, we project habitat suitability changes for the focus species. The models, validated against Lyme disease incidence rates, predicted a 1.5-fold increase in suitable habitats in Latvia opposite to a 4.5-fold decrease in suitable habitats within Ukraine over the coming decades. SHAP values were analyzed to determine the most influential climatic features affecting tick distribution, providing insights for future vector control and disease prevention strategies: here name the drivers in decreasing order. This implies an increased presence of ticks in Scandinavian countries (Sweden, Norway, Finland), Baltic states (Estonia, Latvia, Lithuania), Denmark, and Belarus. These findings largely coincide with our projections regarding bioclimatic suitability for ticks in Ukraine and Latvia. These shifts reflect broader patterns of vector redistribution driven by global warming, with potential expansions in higher altitudes and latitudes. Our findings highlight the urgent need to adapt public health planning to the evolving landscape of vector-borne diseases under climate change.

## 1. Introduction

Climate change is generally considered as one of the main drivers affecting biodiversity [1,2]. A particular interest exists in assessing the potential changes in habitat suitability due to climate change for medically or economically relevant species. For instance, IPCC reports claimed that among other consequences, climate change will change the range of some of the vectors and vector-borne infectious diseases [3, 4]. Tick-borne diseases have been considered a major health problem in recent decades. Ticks classified under the suborder Ixodida are distributed widely in the world and they are very sensitive to climate due their dependence of most of their life cycle stages on a complex combination of climate variables for development and survival [5]. Ticks are vectors for a variety of pathogens, including protozoa, viruses, bacteria and nematodes [6,7], which are considered causative agents of a number of diseases, in particular Lyme borreliosis. In Europe, it is one of the most common tick-borne diseases in humans [8], with *Ixodes ricinus* (Linnaeus, 1758) being the main vector species. The disease has been found to be prevalent in Central Europe where the highest infection rates have been recovered from ixodid ticks [9]. A recent study reported that Central Europe has the highest share of residents with Lyme disease compared to North America [10] (respectively 21% and 9%).

*I. ricinus* is the most abundant and widespread tick species in Europe [11]. It infests a wide variety of wild vertebrate species, as well as other accidental hosts, such as humans, livestock, and pets [12]. In Europe, *I. ricinus* covers a wide geographic range including Scandinavia, the British Isles, Central Europe, France, Spain, Italy, the Balkans, and Eastern Europe, including Ukraine [13] (ECDC 2020). Over the last century, annual mean temperature has risen by 0.7°C globally, and another 1.1°C increase is expected by the end of the 21st century [14]. By affecting vector biology and disease transmission, climate change may have a dual effect on tick occurrence and the spread of tickborne diseases. On one hand, climate change has been shown to drive *I. ricinus* geographical expansion at the northern range margin and/or increase their abundance locally [15-17]. On the other hand, at very high temperatures ticks are vulnerable to desiccation-induced mortality [6]. Furthermore, ticks are less likely to seek hosts when temperatures are high, which may increase tick mortality rates by reducing host-finding success [18]. Therefore, global warming may negatively affect the species’ habitat suitability in the southern parts of its range of distribution [19] and may potentially lead to geographical contractions of its range in areas where the combination of climate variables for development and survival turns out to become less or entirely not favourable. Under these circumstances, there is a need to construct the current climatic niche of *I. ricinus* and project it into future conditions to assess how climatic changes will affect the geographic distribution of the vector. Ultimately, climate-induced changes in vector distribution may affect the epidemiology of vector-borne diseases [20] and are expected to be of profound character [21].

With the global incidence of Lyme disease and other tickborne diseases on the rise, there is a critical need for data-driven tools for assessing the magnitude of this problem and provide scientific-grounded guidelines for public health decision makers. Moreover, there is an interest to identify potential areas of distribution of vector species in order to assess the future infection risks with vector-borne diseases and improve surveillance efforts [22]. In this respect, species distribution modeling (SDM) can be considered as such data-driven numerical tools that combine field observations of species occurrence or abundance with environmental estimates (usually, climatic), which can significantly contribute to predict the potential impacts of climatic changes on species distribution [23]. Modeling the ecological requirements of species to explore future disease transmission patterns has been found challenging [24], however it is essential to assess the potential impacts of climate change on vector distribution, which can provide a theoretical basis for the prevention and control of vector-borne diseases [25,26].

In this study, we used a SDM approach to investigate the distribution of the Lyme disease vector *I. ricinus* under current climate, how climate change predicted by 2030 and 2050 may affect it, and to rank habitat suitability areas for the tick at a country scale with a focus on Ukraine and Latvia. Particularly for Ukraine, the urgency of the problem is exacerbated by the fact that Russia’s armed aggression forced millions of citizens to flee their homes in search of security and temporarily relocated mainly to western and central regions of the country further away from the war zone [27,28]. Therefore, a question raises of how safe are these regions in terms of disease transmission. This could substantially help health officials in managing and monitoring tick distributions, assessing their overlap and potential contact with humans because it is vital to decrease the risk of zoonotic disease transmission.

## 2. Materials and Methods

### 2.1. Tick input data

As input, SDMs require georeferenced biodiversity observations. Faced with limited data for Ukraine, we built species distribution models using European and adjacent occurrences. Presence data was retrieved from online public databases, which were accessed using the R package ‘rgbif’, an interface to the Global Biodiversity Information Facility [29,30] (GBIF.org, 2022), supplemented from the literature [31-34], including Ukrainian and Latvian sources [35-37].

The conducted search for occurrence data yielded a total of 12,095 non-duplicate georeferenced records of *I. ricinus* across Europe and adjacent areas. Using the pre-modeling functions from the ‘flexsdm’ R package, presence records were reduced to 1,827.

### 2.2. Climate data

SDMs are primarily climate-driven, meaning that the variables used to develop them typically portray climatic factors [38,39]. This makes sense because climate is a chief driver of environmental suitability [40]. Information on the bioclimatic parameters was collected as raster layers from three climatic data bases and used separately for building the anticipated SDMs and checking their performances.

From the WorldClim website (http://www.worldclim.com/version2; accessed on 21 November 2022), 19 bioclimatic variables, indicating general trends in precipitation and temperature, including extremes and the seasonality of temperature [41], were downloaded at 2.5’ resolution.

In this study, we used for modeling purposes a set of 16 climatic and 2 topographic variables (the ENVIREM dataset, downloaded from http://envirem.github.io (accessed on 26 November 2022) in a 2.5’ resolution), which were found by recently reconsidering biological significance; many of them are likely to have direct relevance to ecological or physiological processes determining species distributions [42]. These variables are worth consideration in species distribution modeling applications, especially as many of the variables (in particular, potential evapotranspiration) have direct links to processes that are important for species ecology. The included topographic variables are potentially important too, because they can modify the effects of the climate descriptors.

CliMond v.1.2 datasets were downloaded from https://www.climond.org/ at 10’ resolution (accessed on 28 November 2022). These included the core set of 19 bioclimatic variables (temperature and precipitation) and an extended set of 16 additional variables (solar radiation and soil moisture) [43,44].

Climate variables often show high collinearity, and most SDM approaches require the selection of one among strongly correlated variables [45]. In order to carry out such selection, the “removeCollinearity” function in the “virtualspecies” R package was employed [46]. This function analyses the correlation among variables of the provided stack of environmental variables and returns a vector containing names of variables that are not collinear, and also groups variables according to their degrees of collinearity. The collinearity was assessed over the whole extent of Europe and adjacent areas to prevent a collinearity shift when projecting training extents [47]. As climate variables are commonly skewed or have outliers, the Spearman correlation method has been applied using a 0.8 cut-off [48].

Removing high collinearity and the selection of one amongst strongly correlated predictors gave the following results presented in Table 1.

**Table 1.** Results of the selection of uncorrelated predictors (Spearman’s r<0.8)

| Predictor dataset | Selected subsets of predictors |
| --- | --- |
| WorldClim v.2 | Annual Mean Temperature (bio1), Mean Diurnal Range (bio2), Temperature Seasonality (bio4), Maximum Temperature of Warmest Month (bio5), Annual Precipitation (bio12), Precipitation Seasonality (bio15), Precipitation of Driest Quarter (bio17) |
| ENVIREM | Annual PET*, Continentality, Emberger's pluviothermic quotient, mean monthly PET of coldest quarter, mean monthly PET of driest quarter, PET seasonality, mean monthly PET of the wettest quarter, terrain roughness index |
| CliMond v.1.2 | Mean diurnal temperature range (bio2), Minimum temperature of coldest week (bio6), Temperature annual range (bio7), Mean temperature of driest quarter (bio9), Annual precipitation (bio12), Precipitation of driest week (bio14), Precipitation seasonality (bio15), Precipitation of warmest quarter (bio18), Radiation of wettest quarter (bio24), Radiation of warmest quarter (bio26), Radiation of coldest quarter (bio27), Highest weekly moisture index (bio29) |

\*PET = potential evapotranspiration

### 2.3. Modeling procedure

To prepare the input we used pre-modeling functions from the ‘flexsdm’ R package [49]. The calibration area was defined using 500 km buffers around presence points. Filtering the occurrence data was used to reduce sample bias by randomly removing points where they are dense (oversampling) in environmental and geographical space. The modeling functions of the package allow fitting and evaluating different modeling approaches, including individual algorithms, tuned models, and ensemble models. Seven machine learning (ML) SDM methods were employed, including ‘Generalized Additive Models’, ‘Gaussian Process Models’, ‘Generalized Linear Models’, ‘Maximum Entropy’, ‘Artificial Neural Networks’, ‘Random Forest’, and ‘Support Vector Machine’, resulting in a summarizing ensemble model, in which predictions of individual algorithms are combined to produce a consensus distribution, thus reducing model uncertainty and improving model transferability [50].

Models were firstly evaluated using the area under the receiver operating characteristic curve (AUC) [51,52] and the true skill statistic (TSS) [53]. AUC scores range from 0 to 1, with 0 for systematically wrong model predictions and 1 for systematically perfect model predictions; AUC values 0.7 to 0.8 are considered acceptable, values > 0.8 are considered to be good to excellent. TSS values range from -1 to +1, with -1 corresponding to systematically wrong predictions and +1 to systematically correct predictions; TSS values < 0.4 are considered poor, 0.4–0.8 useful, and > 0.8 good to excellent [25]. Because AUC has its drawbacks [54], we also employed the continuous Boyce index provided by the ‘flexsdm’ package, which only requires presences and measures how much model predictions differ from random distribution of the observed presences across the prediction gradients [55]. Thus, it is an appropriate metric in the case of presence-only models. It is continuous and varies between -1 and +1. Positive values indicate a model which present predictions are consistent with the distribution of presences in the evaluation dataset, values close to zero mean that the model is not different from a random model, negative values indicate counter predictions [56].

Secondly, we assumed that maps of habitat suitability of the Lyme disease vector can serve as proxies for the potential distribution of the disease agent itself. Performance of each ensemble model was checked by correlating the obtained habitat suitability values with reported incidence of Lyme disease (calculated as the number of reported cases per 100,000 people) provided by the Johns Hopkins Lyme and Tick-Borne Disease Dashboard (https://www.hopkinslymetracker.org; accessed on 23 December 2022). For this purpose, we used the R-package ‘trafo’ [57] to help select suitable transformations depending on statistical requirements and data being analyzed. To control for spatial non-independence, we used a modified t-test to calculate the statistical significance of the correlation coefficient (a corrected Pearson’s correlation) based on geographically effective degrees of freedom as implemented in the ‘SpatialPack’ package [58].

Eventually, the ‘best’ ensemble SDM was chosen by taking into account the mentioned above performance criteria and on how close is the correlation between *I. ricinus* habitat suitability values and reported incidence of Lyme disease. The best performing SDM was reclassified to areas of low potential habitat suitability, medium and high potential habitat suitability. We defined these thresholds based on Jenks natural breaks, which maximizes the similarity of numbers in groups by minimizing each class average deviation from the class mean, while maximizing each class deviation from the means of the other groups. The Jenks natural break provides a uniform interface to finding class intervals for continuous numerical variables [59].

### 2.4. Feature importance

A critical challenge in the adoption of ML models is their inherent lack of interpretability, often referred to as the “black box” problem. Shapley Additive exPlanations (SHAP) [60-62] represents a significant advancement in addressing this concern by providing a framework for understanding how ML models arrive at their predictions. SHAP offers various advantages over other feature importance methods, including its model-agnostic nature. Leveraging game theory concepts, SHAP provides a robust framework for feature importance attribution in ML models, despite the number of used variables. The SHAP value quantifies the magnitude and direction (positive or negative) of the feature’s influence on the prediction [60-61]. In our case the relative importance of predictors was measured with mean absolute Shapley values, which can be interpreted as the magnitude of the relative contribution of a predictor (feature) towards a model output [63]. The R package ‘shap-values’ (https://github.com/pablo14/; author Pablo Casas; accessed 10 December 2023) in a modified version was used to calculate SHAP values for a selected model. A summary plot offers a comprehensive view of the most influential features in a model and ranks features based on their effect on the model’s predictions. In other words, SHAP values can be integrated into global explanations such as variable importance, but using a completely different from various widely applied so far feature selection methods [64]. Until only now, the use of SHAP for explaining which environmental factors (including climatic) influence the spatial distribution of species occurrences has started to be explored more widely [65-70].

Maps of habitat suitability in the GeoTIFF format were processed and visualized in SAGA GIS [71], statistical data was analyzed using the PAST software package [72,73] and/or the R environment [74].

## 3. Results

As a result of the modeling, we obtained models with a sufficiently high evaluation. From figures in Table 2 it is evident that SDMs built using various selected subsets of predictors from the WorldClim v.2, ENVIREM and CliMond v.1.2 datasets performed satisfyingly, with AUC values considered mostly to correspond to good and excellent, and useful TSS values. However, more weight in this respect should be given to the continuous Boyce index, which is considered one of the most appropriate metrics for assessing model predictions applied to presence-only datasets and a more reliable metric than AUC [75,76]. Accordingly, values reported in Table 1 indicate models in which present predictions are highly consistent (>0.9) with the distribution of presences in the datasets. Also, the continuous Boyce index shows a high uniformity of performance of the obtained SDMs and where now it reveals a challenge to picking the ‘best’ model.

**Table 2.** Evaluation metrics for SDMs built using various selected subsets of predictors.

| Predictor dataset/<br>subset | Evaluation metrics |  |  |  |  |  |
| --- | --- | --- | --- | --- | --- | --- |
|  | Area Under<br>Curve (AUC) | SD* of<br>AUC | True Skill<br>Statistic<br>(TSS) | SD of<br>TSS | Continuous<br>Boyce index<br>(BOYCE) | SD of<br>BOYCE |
| WorldClim v.2 | 0.84 | 0.01 | 0.59 | 0.02 | 0.91 | 0.07 |
| ENVIREM | 0.84 | 0.01 | 0.56 | 0.02 | 0.92 | 0.06 |
| CliMond v.1.2<br>current | 0.79 | 0.06 | 0.50 | 0.12 | 0.91 | 0.07 |
| CliMond v.1.2<br>A1B scenario for<br>2030 | 0.76 | 0.06 | 0.46 | 0.13 | 0.87 | 0.18 |
| CliMond v.1.2<br>A1B scenario for<br>2050 | 0.74 | 0.08 | 0.41 | 0.12 | 0.69 | 0.16 |

\*SD = standard deviation

The ‘flexsdm’ package supports model evaluation based on a number of mentioned above performance metrics. Amongst the threshold criteria maximum training sensitivity plus specificity was considered a reasonable choise [77,78].

Further, the ensemble models for current and projected climates were clipped to Ukraine and Latvia and classified using the corresponding module in SAGA GIS in accordance with Jenks natural breaks into three categories, i.e., “high”, “medium” and “low”, habitat suitability (Figs. 1 - 3).

**Figure 1.**
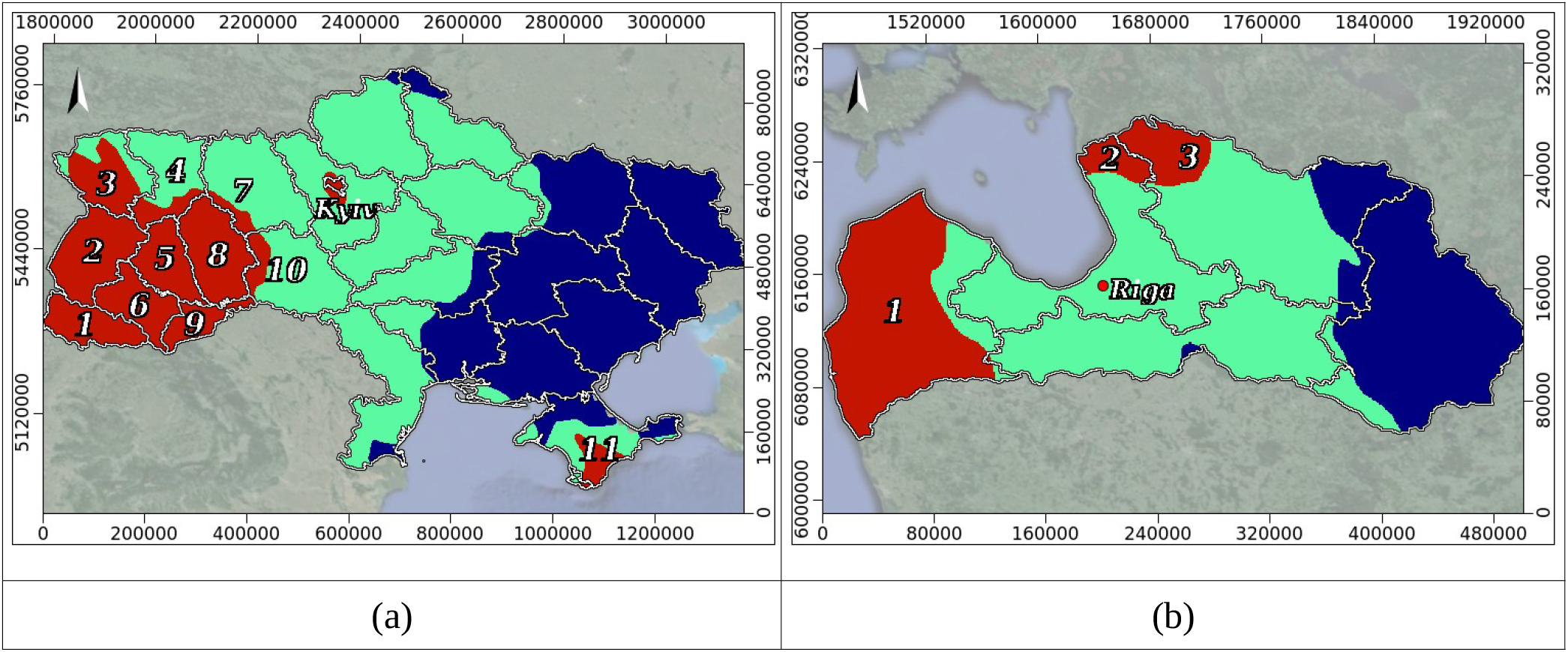
Jenks natural breaks maps of habitat suitability (HS) for *Ixodes ricinus* under current bioclimatic conditions: (**a**) Ukraine: red, green and blue respectively for HS of high (0.52 - 0.71), medium (0.29 - 0.52), and low (0.08 - 0.29) value (Oblasts: 1 - Zakarpatska, 2 - Lvivska, 3 - Volynska, 4 - Rivnenska, 5 - Ternopilska, 6 - Ivano-Frankivska, 7 - Zhytomyrska, 8 - Khmelnytska, 9 - Chernivetska, 10 - Vinnytska, 11 – Crimea); (**b**) Latvia: red, green and blue respectively for HS of high (0.62 - 0.77), medium (0.46 - 0.62), and low (0.32 - 0.46) values (Regions: 1-Kurzeme, 2 - Pieriga, 3 – Vidzeme). Administrative units according to OECD, 2022.

Upon analyzing the models, we can predict that in the north and south of its range, the potential distribution of *Ixodes ricinus* may take different directions: for instance in Ukraine, in the south of its range, predicted increased fragmentation may lead to a significant decrease in tick populations. On the contrary, in the northern part of Ukraine, the models predict an increase in the potential suitable territory for the spread of the species.

## 4. Discussion

As already implied, there may be another approach, tentatively tagged “biological”, to finding a model that would meet our objective aimed to enhance health officials in managing and monitoring tick and tick-borne disease distributions. We assume this can be accomplished by correlating the obtained habitat suitability values of each SDM with reported incidence of Lyme disease and hypothesizing that a closer correlation would point towards a model that fits our needs the most. Existing studies in Europe mainly focus on acarological risk assessment, with very limited investigations exploring Lyme disease occurrence in human [79]. As an example, incidences of Lyme disease in Ukraine are scarcely reported so that proper statistical analyses cannot be achieved. Therefore, we used data from Romania since this neighbouring country presents fairly similar climate compared to Ukraine, especially if accounting for warm season temperatures when tick activity is the highest [80] (https://www.worlddata.info/, accessed on 10 January 2023). From the Johns Hopkins Lyme and Tick-Borne Disease Dashboard annual incidences of Lyme disease were downloaded for 41 counties and Bucharest Municipality for a 10 year period (from 2012 to 2021) and averaged. Corresponding averaged habitat suitability values for each administrative entity, represented by a polygon shapefile, were obtained using the ‘grid statistics for polygons’ module in SAGA GIS. Firstly, using the R package ‘trafo’ suitable transformations depending on statistical requirements and the data being analyzed were assessed by checking assumptions of normality, homoscedasticity and linearity. By log-transforming the annual incidences of Lyme disease using the formula ln(x+1), all specified assumptions were shown to be acceptable (p>0.05). Secondly, using log-transformed annual incidences of Lyme disease and the averaged habitat suitability values regarding the considered administrative units in Romania, corrected Pearson’s correlation coefficient (r) controlling for spatial non-independence were calculated. Results concerning each of the SDMs reveal the following: for data extracted from the SDM based on the WorldClim v.2 subset r=0.3815, p-value=0.0584, meaning no confirmation of statistical significance; in the case of the SDM based on the ENVIREM subset r=0.3989, the p-value being 0.0422, and in the case of the SDM based on the CliMond v.1.2 subset r=0.59150, with a p-value of 0.0015. For the last two subsequent cases conclusions regarding correlations are of sound statistical significance, however a stronger correlation coefficient is found for the SDM built on the CliMond v.1.2 subset, therefore we can conclude that this particular model is best fit for our purpose.

In this study, besides the investigation of present distribution of the tick *I. ricinus*, we aimed at assessing how future climatic changes predicted for 2030 and 2050 could affect it in order to be prepared for proactive management actions involving both the vector and spread of Lyme disease. In this place we employed the CliMond v.1.2 data set projection of future climate for 2030 and 2050 generated for the A1B scenario for emissions of greenhouse gases and sulphate aerosols [37], which envisages a balanced emphasis on all energy sources. The produced by using the ‘flexsdm’ algorithm ensemble models performed fairly good (see Table 2), with an acceptable AUC value, useful TSS, and high value of the continuous Boyce index. Somewhat poorer was the performance of the 2050 model, but yet in acceptable limits.

In terms of the current bioclimate, areas of high habitat suitability for *I. ricinus* in Ukraine are predicted to occupy around 20% of the country, with locations found predominantly in the west (Figure 1a). Other smaller patches of habitat suitability were found around Kyiv and in the south of Crimea, where tick are the most abundant representative of the family [81]. With regard to health issues, as far as since the start of the full-scale Russian invasion in Feb, 24, 2022 over 12 million people have fled their home in Ukraine while more than 5 additional million people are estimated to be displaced inside Ukraine (The UN Refugee Agency; https://www.unhcr.org/emergencies/ukraine-emergency), such prediction may have potential consequences. Further while refugee movements are associated with an increase in infectious disease transmission and are likely to affect zoonotic disease risks, it yet remains unclear how forced migrations affect disease dynamics [82]. Human susceptibility to disease during forced migration might increase due to exhaustion, malnutrition and stress arising from displacement, magnified by crowded and substandard living conditions [72,73]. As for Latvia, under current bioclimatic conditions, areas of high habitat suitability for the tick occur on 22% of the territory of the country, mainly in the west and to a lesser extent in the north (Figure 1b).

By 2030, in Ukraine, our models predict an around two-fold reduction of areas of high habitat suitability for *I. ricinus*, to <10% of the country (Figure 2a). These are expected to be fully located in the west, whereas patches of habitat suitability around Kyiv and in the south of Crimea most likely will disappear. By 2050, in Ukraine, highly suitable habitats for *I. ricinus* are expected to continue their contraction (to 4.5% of the country), with potential reduction to Lvivska and Ivano-Frankivska oblasts, despite a patch of high habitat suitability may appear in the southern region of Odesa (Figure 3a). On the contrary, in Latvia, areas of high habitat suitability for *I. ricinus* will possibly expand and occupy up to 27% of the territory of the country by 2030, and up to 33% in 2050, engulfing the entire western region of Kurzeme (Figure 2b).

**Figure 2.**
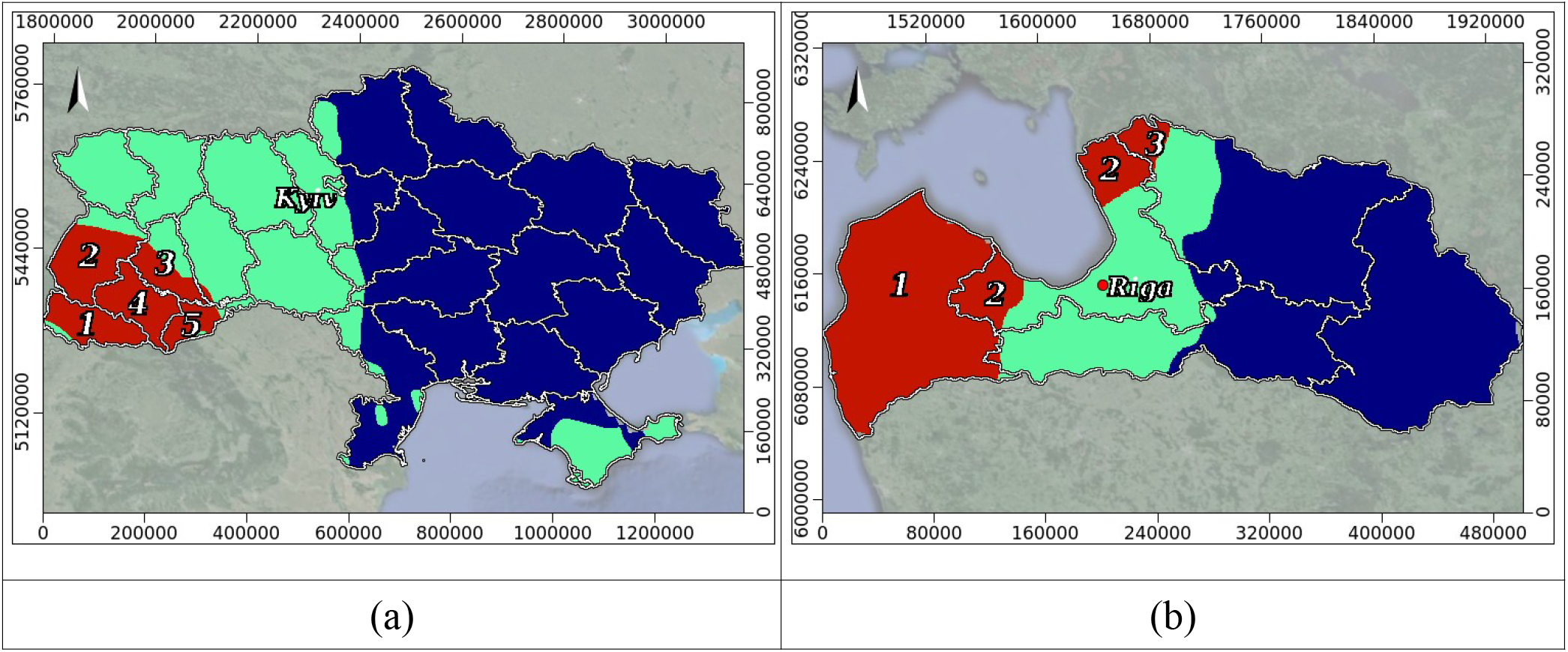
Jenks natural breaks maps of habitat suitability (HS) for *Ixodes ricinus* as predicted by our models by 2030: (**a**) Ukraine: red, green and blue correspondingly denote areas with high (0.61-0.77), medium (0.36-0.61), and low (0.1-0.36) HS. Oblasts: 1 - Zakarpatska, 2 - Lvivska, 3 - Ternopilska, 4 - Ivano-Frankivska, 5 - Chernivetska; (**b**) Latvia: red, green and blue correspondingly denote areas with high (0.62-0.76), medium (0.48-0.62), and low (0.34-0.48) HS. Regions: 1-Kurzeme, 2 - Pieriga, 3 - Vidzeme.

**Figure 3.**
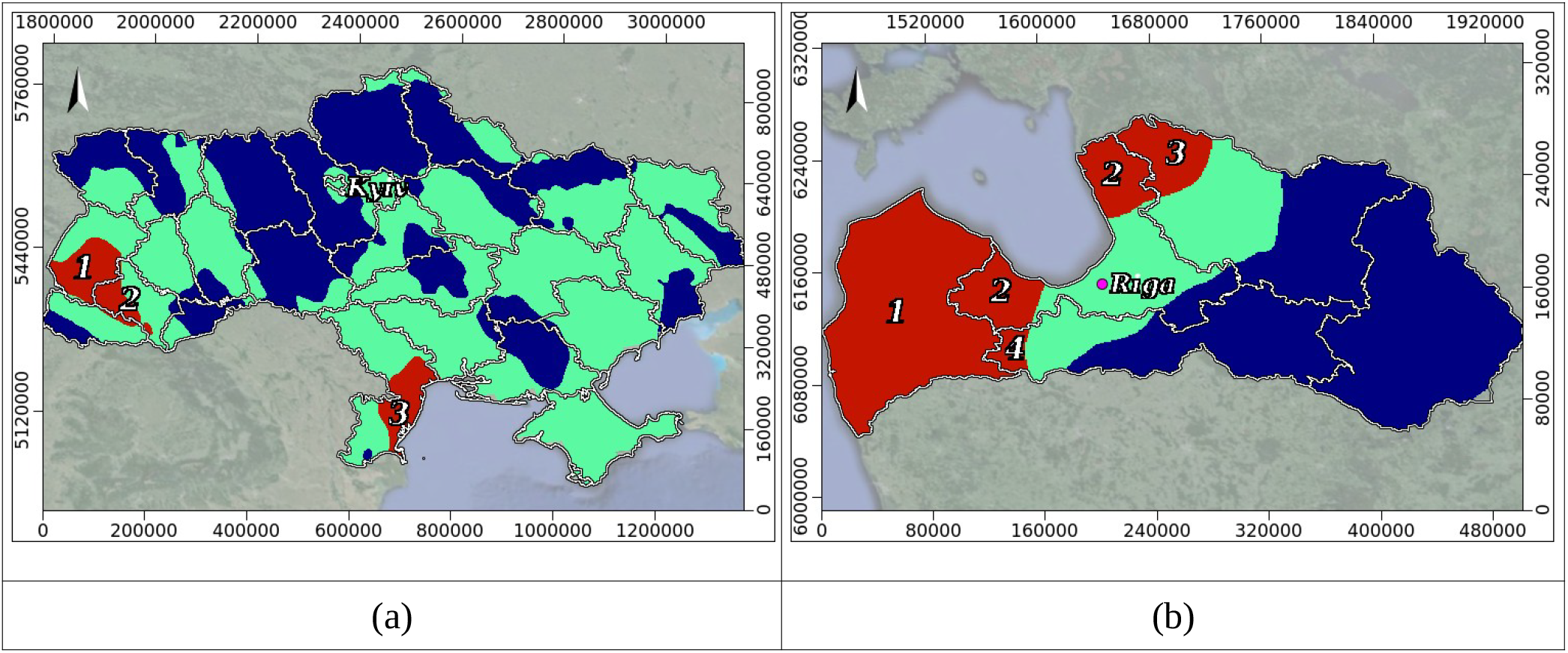
Jenks natural breaks maps of habitat suitability (HS) for *Ixodes ricinus* in different countries under future climate for 2050: (**a**) Ukraine: red, green and navy blue correspondingly denote areas with high (0.63-0.73), medium (0.45-0.63), and low (0.33-0.45) HS. Oblasts: 1 - Lvivska, 2 - Ivano-Frankivska, 3 – Odeska; (**b**) Latvia: red, green and navy blue correspondingly denote areas with high (0.69 - 0.8), medium (0.51 - 0.69), and low (0.33 - 0.51) HS. Regions: 1-Kurzeme, 2 - Pieriga, 3 - Vidzeme, 4 - Zemgale.

Our results regarding prospects of bioclimatic suitability for the castor bean tick in Ukraine and Latvia are consistent with findings from previous studies. Indeed, previously published SDMs have predicted that Ixodes ticks will increase their distribution and abundance under global warming, the most generally envisaged scenario being an expansion of geographical range for ticks, including *I. ricinus*, with potential increases in abundance in some regions [83-85]. Bioclimatic models also predict a shift in the distribution of the tick species particularly at higher altitudes and latitudes, like Scandinavia, the Baltic states, and Belarus. Conversely, tick populations are expected to decline in areas like the Alps, Pyrenees, central and western Italy, and northwestern Poland. This means an increase in tick presence for Scandinavian countries (Sweden, Norway, Finland), the Baltic states (Estonia, Latvia, Lithuania), Denmark, and Belarus [86].

In our study, we analyzed SHAP values to understand feature importance governing current tick distribution in Ukraine and Latvia by calculating the average absolute values for each feature. This permits to provide an idea of which features are generally most important in influencing predictions within the area of our countries of interest (Figure 4). Given their extended off-host existence as generalist parasites, ixodid ticks, including the castor bean tick, exhibit a marked sensitivity to both temperature and humidity [87]. This sensitivity arises from the critical influence these environmental factors exert on physiological processes and desiccation rates, both of which are essential for tick survival and life cycle completion [33,88]. In our study, SHAP-analysis suggests that rainfalls during the warmest quarter (Bio18 in Figure 4a) has a particularly strong influence on predictions for Ukraine. Checking the correlation between Bio18 and the ensemble SDM for current climate demonstrated a strong relationship too, that could be explained by the fact that ‘precipitation during the warmest quarter’ acts as a trade-off factor for other environmental variables, including temperature and humidity. Overall, the optimal bioclimatic environment for the tick species seems to be an intricate balance of moderate temperatures, high humidity, and sufficient rainfall to maintain that humidity, despite the fact that the correlation between humidity and precipitation is not straightforward [89-91].

**Figure 4.**
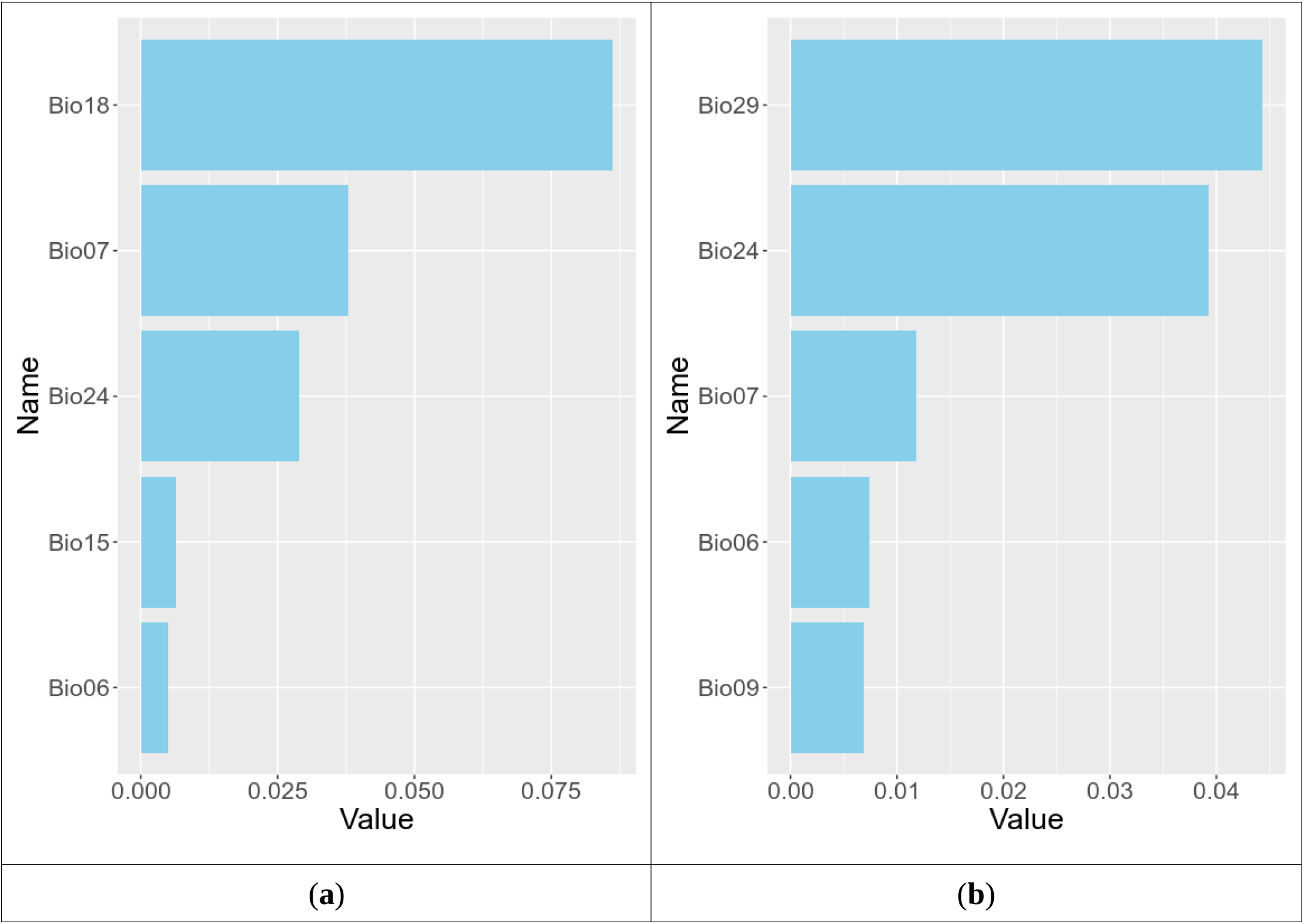
Absolute summary plots of the datasets for Ukraine (**a**) and Latvia (**b**), where the average absolute value of the SHAP values for each variable is taken in order to obtain a bar chart as a function of the contribution of each variable to the prediction of the CliMond v.1.2 current model. The variables are ordered from most (top) to least (bottom) important. The y-axis represents the variables used in the study, which refer to: Bio6 - minimum temperature of coldest week, Bio7 - temperature annual range, Bio9 - mean temperature of driest quarter, Bio15 - precipitation seasonality, Bio18 - precipitation of warmest quarter, Bio24 - radiation of wettest quarter, Bio29 - highest weekly moisture index. The x-axis represents the corresponding SHAP value.

In addition to climate, the distribution of *I. ricinus* may also be affected by factors such as land cover, in particular vegetation. Ticks require moisture and humidity to survive and dense vegetation provides these conditions by shading the ground and preventing moisture loss [92]. Additionally, some stages of the tick lifecycle feed on the blood of various animals that frequent forested and other vegetated areas [12]. Because in our modeling we focused on bioclimatic predictors, strictly speaking vegetation features were not used, however we tested the connection between Bio18 and NDVI (Normalized Difference Vegetation Index). NDVI is a measure of vegetation health and density. Corresponding rasters for the warmest quarter (June - August) were downloaded from the EDIT geoplatform (http://edit.csic.es/GISdownloads.html). Previous studies have shown that areas with higher NDVI tend to be more suitable habitat for *I. ricinus* ticks [93]. Indeed, we found fairly strong correlations for summer months (June, July, and August). In our opinion, once again this highlights the importance of the ‘precipitation of warmest quarter’ factor and/or associated with it factors in shaping the tick’s ecological niche and distribution, specifically in Ukraine.

In fact, for Latvia we have an analogous situation, where the most important features in influencing predictions within the country are suggested to be Bio29 (‘highest weekly moisture index’) and Bio24 (‘radiation of wettest quarter’) (Figure 4b). Both characterize the moisture regime making a direct or indirect emphasis on the warm time of the year. The study area experiences its highest weekly moisture index in July. Secondly, the wettest quarter occurs from July to September, with peak rainfall in July.

## 5. Conclusions

As a result of our study, global warming is one of the most important factors in redistribution of the tick populations and the outbreak of tick-borne diseases and support recent studies [94,95]. SDMs built using various selected subsets of predictors from the WorldClim v.2, ENVIREM and CliMond v.1.2 datasets performed satisfyingly, particularly the continuous Boyce index shows a high uniformity of performance of the obtained SDMs. In another approach by correlating the obtained habitat suitability values of each SDM with reported incidence of Lyme disease we found a stronger correlation coefficient for the SDM built on the CliMond v.1.2 subset, therefore we concluded that this particular model is best fit for our purpose.

SDMs employing the CliMond v.1.2 data set projection for future climates (2030 and 2050) generated for the A1B scenario for emissions of greenhouse gases and sulphate aerosols performed fairly good, somewhat poorer was the performance of the 2050 model, but yet in acceptable limits. In terms of the current bioclimate, areas of high habitat suitability for *I. ricinus* in Ukraine are predicted to occupy around one fifth of the country, with locations found predominantly in the west thus highlighting here the elevated risk of disease transmission. Regarding Latvia, areas of high habitat suitability for the tick are predicted to occur mainly in the west and to a lesser extent in the north of the country. Modeling suggests a significant reduction of highly suitable habitat for *I. ricinus* within Ukraine over the coming decades, whereas for Latvia expansion of such suitable habitat is expected to continue, accordingly aggravating future health concerns.

We consider the relative importance of predictors in our case study was successfully measured with mean absolute Shapley values, underlining the pivotal significance of moisture conditions in the warm months of the year for creating highly suitable habitat for the tick species.

## Author Contributions

Conceptualization, V.T., I.K., M.P. and O.N.; methodology, V.T. and O.N.; software, V.T. and O.N.; validation, V.T. and O.N.; formal analysis, V.T. and O.N.; investigation, V.T. O.N., M.P., and A.Č.; resources, V.T., I.K., O.N. and M.P.; data curation, V.T., I.K., O.N., M.P., and A.Š.; writing—original draft preparation, V.T. and O.N.; writing —review and editing, V.T., O.N., I.K., M.P., A.Č., and A.Š.; visualization, V.T.; supervision, V.T., O.N., I.K., M.P., A.Š.; project administration, V.T., O.N., M.P., and A.Š. All authors have read and agreed to the published version of the manuscript.

## Data Availability Statement

Data derived from public domain resources:

1. Species presence records *Ixodes ricinus* (Linnaeus, 1758):
  - GBIF.org (22 May 2024) GBIF Occurrence Download https://doi.org/10.15468/dl.bf6zjp
2. Environmental data:
  - The WorldClim v.2 website (http://www.worldclim.com/version2; accessed on 21 November 2022).
  - The ENVIREM dataset downloaded from (http://envirem.github.io; accessed 26 November 2022).
  - The CliMond v.1.2 (current, 2030, 2050) datasets were downloaded from https://www.climond.org/ at 10’ resolution (accessed on 28 November 2022).
3. Software:
  - Maxent, https://biodiversityinformatics.amnh.org/open_source/maxent/
  - The R package ‘rgbif’, https://github.com/ropensci/rgbif
  - The R package ‘flexsdm’, https://sjevelazco.github.io/flexsdm/, https://besjournals.onlinelibrary.wiley.com/doi/full/10.1111/2041-210X.13874
  - The R package “virtualspecies”, https://borisleroy.com/virtualspecies/ https://cran.r-project.org/web/packages/trafo/index.html
  - The R package ‘shap-values’, https://github.com/pablo14/
  - The R-package ‘trafo’, https://github.com/cran/trafo
  - The R-package ‘SpatialPack’, http://spatialpack.mat.utfsm.cl/
  - The PAST software package and/or the R environment [72,73,74], https://www.nhm.uio.no/english/research/resources/past/
  - The Saga GIS software, https://saga-gis.sourceforge.io/en/index.html

## Acknowledgments

We acknowledge the Emys-R project (https://emysr.cnrs.fr) under the Biodiversa+ and Water JPI joint call for research projects, under the BiodivRestore ERA-NET Cofund (GA N°101003777), with the EU and the funding organizations Agence Nationale de la Recherche (ANR, France, grant ANR-21-BIRE-0005), Bundesministerium für Bildung und Forschung (BMBF, Germany, grant 16LW015), the State Education Development Agency (VIAA, Latvia, grant ES RTD/2022/2), and the National Science Center (NSC, Poland, grant 2021/03/Y/NZ8/00101). We are also thankful for the support from the Collège de France and Agence Nationale de la Recherche (ANR) through the PAUSE ANR Ukraine program (Nekrasova O., grant ANR-23-PAUK-0074), and the project “Ecological and socioeconomic thresholds as a basis for defining adaptive management triggers in Latvian pond aquaculture” (lzp-2021/1-0247), project 16-00-F02201-000002 for the use of the mobile complex of scientific laboratories for research purposes.

## Disclaimer/Publisher’s Note

The statements, opinions and data contained in all publications are solely those of the individual author(s) and contributor(s) and not of MDPI and/or the editor(s). MDPI and/or the editor(s) disclaim responsibility for any injury to people or property resulting from any ideas, methods, instructions or products

